# Mutation of Arabidopsis SME1 and Sm core assembly improves oxidative stress resilience

**DOI:** 10.1101/2022.04.12.488072

**Authors:** Patrick Willems, Valerie Van Ruyskensvelde, Takanori Maruta, Robin Pottie, Álvaro Daniel Fernández-Fernández, Jarne Pauwels, Matthew A. Hannah, Kris Gevaert, Frank Van Breusegem, Katrien Van der Kelen

## Abstract

Alternative splicing is a key posttranscriptional gene regulatory process, acting in diverse adaptive and basal plant processes. Splicing of precursor-messenger RNA (pre-mRNA) is catalyzed by a dynamic ribonucleoprotein complex, designated the spliceosome. In a suppressor screen, we identified a nonsense mutation in the Sm protein SME1 to alleviate photorespiratory H_2_O_2_-dependent cell death in catalase deficient plants. Similar attenuation of cell death was observed upon chemical inhibition of the spliceosome, suggesting pre-mRNA splicing inhibition to be responsible for the observed cell death alleviation. Furthermore, the *sme1-2* mutants showed increased tolerance to the reactive oxygen species inducing herbicide methyl viologen. Both an mRNA-seq and shotgun proteomic analysis in *sme1-2* mutants displayed a constitutive molecular stress response, together with extensive alterations in pre-mRNA splicing of transcripts encoding metabolic enzymes and RNA binding proteins, even under unstressed conditions. Using SME1 as a bait to identify protein interactors, we provide experimental evidence for almost 50 homologs of mammalian spliceosome-associated protein to reside in the *Arabidopsis thaliana* spliceosome complexes and propose roles in pre-mRNA splicing for four uncharacterized plant proteins. Furthermore, like in *sme1-2*, a mutant in the Sm core assembly protein ICLN resulted in a decreased sensitivity to methyl viologen. Taken together, these data show that both a perturbed Sm core composition and assembly results in the activation of a defense response and enhanced resilience to oxidative stress.

## Introduction

Alternative splicing (AS) increases gene expression complexity by producing diverse protein coding transcripts from a single precursor-messenger RNA (pre-mRNA). Studies exploring transcript diversity in *Arabidopsis thaliana* (Arabidopsis) estimate approximately 60% of protein-coding genes to be alternatively spliced (Marquez et al., 2012; Cheng et al., 2017; Zhang et al., 2017; Laloum et al., 2018). Pre-mRNA splicing is catalyzed by dynamic macromolecular ribonucleoprotein (RNP) complexes, together constituting the spliceosome (Kambach et al., 1999; Matera and Wang, 2014). The major spliceosome consists of five RNA-protein complexes, designated the U1, U2, U4, U5, and U6 small nuclear ribonucleoproteins (snRNPs) that bind uridine-rich small nuclear RNA (snRNA) and include Smith antigen (Sm) proteins and other associated proteins (Lee and Rio, 2015). The seven evolutionary conserved Sm proteins (B, D1, D2, D3, E, F, and G) are surrounding the snRNA in a heptameric ring-shaped complex via a bipartite Sm sequence motif (Kambach et al., 1999; Salgado-Garrido et al., 1999; Veretnik et al., 2009). The Sm ring and snRNPs are assembled in a multi-step pathway mediated by the survival motor neuron (SMN) complex (Li et al., 2014). Next to Sm proteins, eight ‘Like-Sm’ (LSm) proteins form two different heptameric complexes: the LSm2-8 complex, a component of the U6 snRNP performing pre-mRNA splicing (Achsel et al., 1999;Verdone et al., 2004), and the LSm1-7 complex involved in mRNA decapping (Tharun et al., 2000).

In plants, AS is an important posttranscriptional regulation mechanism in response to environmental perturbations (Laloum et al., 2018). Accordingly, thousands of pre-mRNAs of genes are alternatively spliced during cold acclimation (Calixto et al., 2018), salt stress (Feng et al., 2015), drought and heat stresses (Liu et al., 2018). Concomitantly, Arabidopsis splicing-related proteins themselves regulate pre-mRNA splicing activity in a stress dependent manner (Feng et al., 2015; Perea-Resa et al., 2016; Carrasco-López et al., 2017; Laloum et al., 2018). For instance, LSm8 mutation caused differential splicing under stress, correlating with improved cold and hypersensitivity to salt stress, thereby indicating an environment dependent regulation by the LSm2-8 complex (Carrasco-López et al., 2017). Besides the LSm2-8 complex, also Sm proteins and proteins facilitating snRNP assembly have been associated with stress responses. A mutation of SmD3-b and the snRNP assembly proteins PRMT5 and ICLN results in increased resistance to a virulent oomycete in Arabidopsis (Huang et al., 2016). And, more recently, the Sm protein SME1 was associated with low temperature acclimation, with loss-of-function mutants displaying developmental defects at lower temperatures (Capovilla et al., 2018;Huertas et al., 2019).

So far, our lab identified in a second-site suppressor screen mutations of the metabolic enzyme GLYCOLATE OXIDASE2 and the transcription factor SHORT-ROOT to alleviate photorespiratory cell death in catalase (*cat2-2*) deficient plants (Kerchev et al., 2016;Waszczak et al., 2016). Under photorespiration-promoting growth conditions, Arabidopsis mutants deficient in the hydrogen peroxide (H_2_O_2_)-scavenging CATALASE2 (*cat2-2*) display a loss of PSII maximum efficiency (F_v_’/F_m_’) and develop cell death lesions (Vanderauwera et al., 2011;Kerchev et al., 2015). Here, we describe a nonsense mutation in *SME1* that suppresses *cat2-2* dependent cell death and exhibits a constitutively activated stress response associated with an enhanced tolerance to the ROS stimulating herbicide methyl viologen. mRNA-seq analysis of *sme1-2* mutants revealed pre-mRNA splicing alterations, of which intron retention was the predominant AS event. In a tandem affinity purification (TAP) experiment using SME1 as a bait, almost 50 characterized or hypothesized components of the plant spliceosome were detected. Similarly to *SME1* mutation, mutation of one of its interactors ICLN, involved in snRNP assembly, resulted in stunted growth under low temperatures and enhanced tolerance to methyl viologen. Taken together, our results suggest that perturbation of the Sm protein SME1 or snRNP assembly triggers a constitutive stress response enabling resilience to oxidative stress.

## RESULTS

### *SME1* mutation alleviates *cat2-2* photorespiratory cell death

Previously, we reported an EMS-based genetic suppressor screen to identify second-site mutations that attenuate an H_2_O_2_-dependent cell death phenotype in *cat2-2* mutants (Kerchev et al., 2016;Waszczak et al., 2016). Here, we describe mutant *40*.*2* that displayed a prominent alleviated cell death phenotype, together with a partial restoration of photosystem II maximum efficiency (F_v_’/F_m_’; Figure 1A and 1B). We mapped a recessive causative non-sense mutation in the second exon (‘CAA’ [Gln22] to ‘TAA’→*) of the *SME1* gene (AT2G18740), resulting in a premature stop codon that truncates 67 AA C-terminal residues of the 88 AA full-length protein (Figure 1C and Supplemental Figure 1). To confirm the *SME1* mutation as the causative mutation in *40*.*2*, we crossed *cat2-2* with a homozygous T-DNA insertion allele *sme1-2* (Huertas et al., 2019; also designated *pcp-1* in Capovilla et al., 2018). The T-DNA insertion resides in close proximity to the *40*.*2* nonsense mutation (Figure 1C) and the *cat2-2 sme1-2* double mutant showed a nearly identical mitigation of the *cat2-2* F_v_’/F_m_’ decrease and cell death phenotype (Figure 1A and 1B), confirming that a mutation in *SME1* alleviates H_2_O_2_-induced cell death in *cat2-2*. In addition, *40*.*2* rosette size was reduced (Figure 1B and 1D) as reported before for *sme1-2* (Capovilla et al., 2018; Huertas et al., 2019). When grown in soil at 21°C in long day conditions under a moderate light regime (200 µmol.m^-2^.s^-1^), *cat2-2* plants triggered biotic defense responses and developed hypersensitive response (HR)-like lesions (Chaouch et al., 2010) covering on average 7.3% of the leaf area (Figure 1D and 1E). In contrast, wild-type and *sme1-2* plants presented no HR-like lesions, while lesion formation was mitigated in the *40*.*2* mutant (2.6% leaf area) and nearly abolished in the *cat2-2 sme1-2* double mutant (Figure 1E).

**Figure 1.**
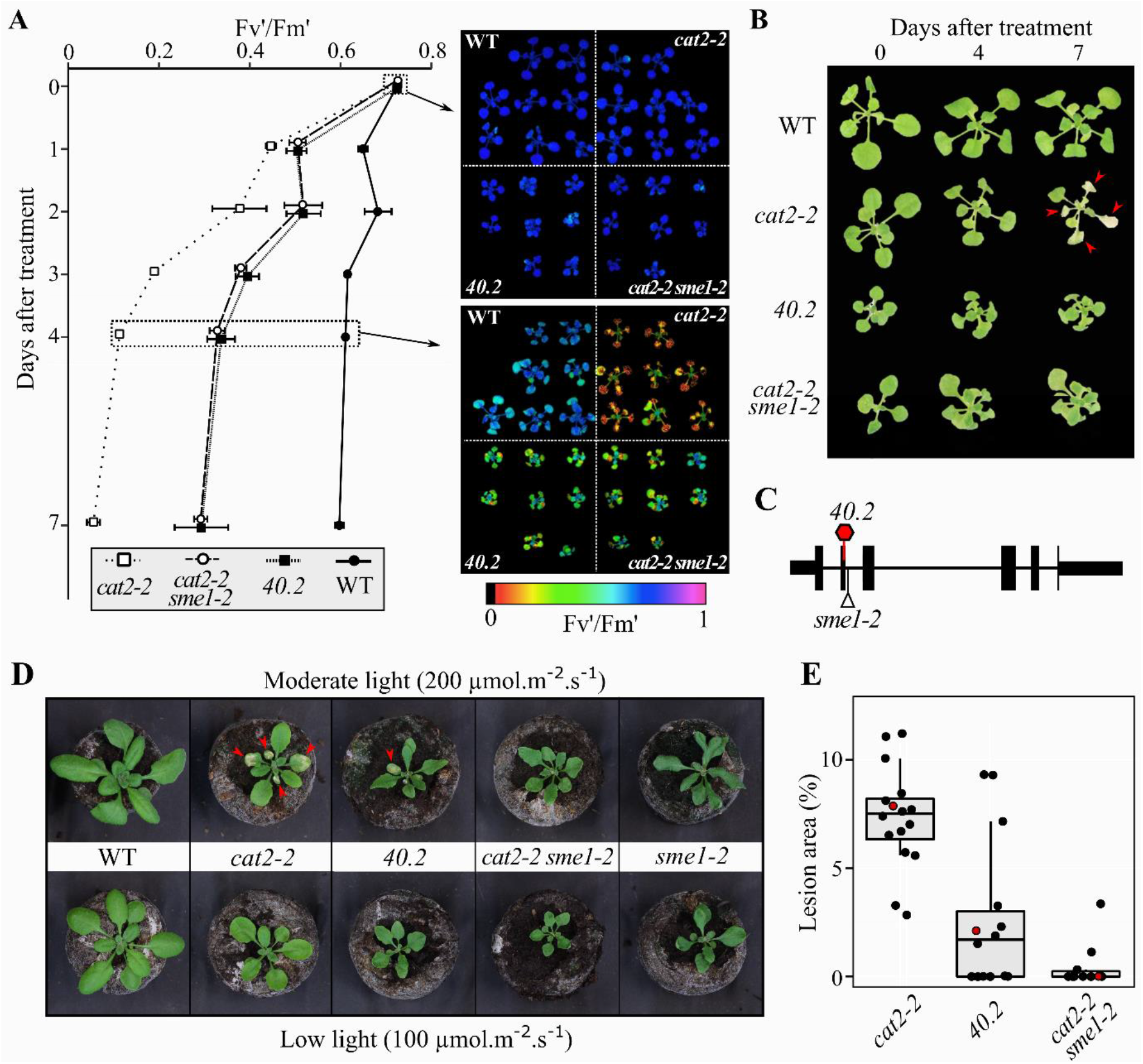
*SME1* mutation mitigates H_2_O_2_-induced cell death in *cat2-2*. A, (*Left*) Decrease in PSII maximum efficiency (F_v_’/F_m_’) in 3-week-old seedlings (n = 3, mean ± SE) under photorespiratory stress conditions caused by restricted gas exchange-continuous light (RGCL) regime (*Right*). Representative color-coded images of F_v_’/F_m_’ taken before (t = 0) and after 4 days of RGCL treatment. B, Representative images of seedlings before (t = 0) and after 4 and 7 days of RGCL treatment. Arrows indicate foliar lesion formation. C, Gene diagram showing the *sme1-2* T-DNA insertion alleles (SALK_089521) and the *40*.*2* nonsense mutation (red hexagon, see also Supplemental Figure 1). Exons and introns are displayed as thick and thin lines, respectively, and UTRs as middle-sized lines. D, Representative pictures of 3-week-old plant grown under moderate (200 µmol.m^-2^.s^-1^) and low light (100 µmol.m^-2^.s^-1^) in a long-day photoperiod (16 h light/8 h dark). Arrows indicate foliar lesion formation. E, Boxplot of lesion leaf area percentage for *cat2-2, 40*.*2* and *cat2-2 sme1-2* when grown under moderate (200 µmol.m^-2^.s^-1^) light in long-day photoperiod. Red data points indicate plants used for representative pictures in panel D.

As it was previously shown that *SME1* mutation perturbs pre-mRNA splicing (Capovilla et al., 2018; Huertas et al., 2019), we tested whether chemical perturbation of pre-mRNA splicing resulted in a similar protection against H_2_O_2_-induced cell death. The polyketide herboxidiene (GEX1A) is a cancer drug targeting the spliceosome U2 snRNP-associated factor 155 and thereby inhibits pre-mRNA splicing (Hasegawa et al., 2011). In fact, it was originally described as a herbicide (Miller-Wideman et al., 1992) and causes pre-mRNA splicing defects triggering stress signaling pathways (AlShareef et al., 2017). Here, we transferred one-week-old *sme1-2, cat2-2 sme1-2, cat2-2*, and wild-type seedlings to growth medium supplemented with 0.1 µM and 0.2 µM GEX1A (Figure 2A).

**Figure 2.**
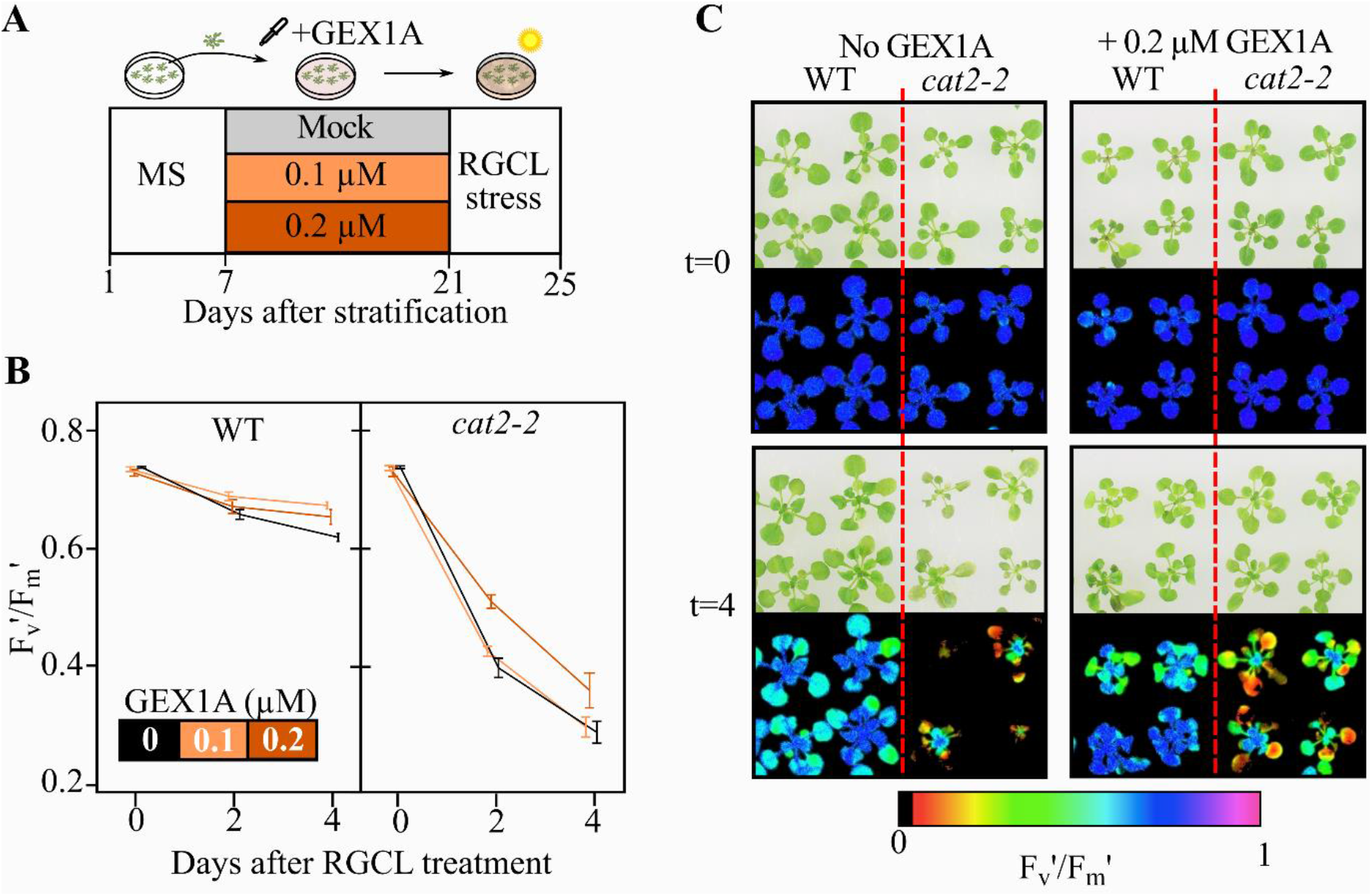
Chemical spliceosome inhibition improves tolerance to photorespiratory stress. A, Experimental set-up of GEX1A treatment. B, PSII maximum efficiency (F_v_’/F_m_’) in 3-week-old seedlings (n = 4 and 2 for stressed and unstressed seedlings, respectively, mean ± SE) under photorespiratory stress conditions caused by restricted gas exchange-continuous light (RGCL) regime. Data are color-coded according to GEX1A concentration. See Supplemental Figure 2A-B for growth parameters and F_v_’/F_m_’ decrease in *sme1-2* and *cat2-2 sme1-2*. C, Representative bright-field and color-coded pictures of F_v_’/F_m_’ of wild-type (WT) and *cat2-2* seedlings taken before (t = 0) and after 4 days (t = 4) of RGCL treatment in control media (*Left*) and media supplemented with 0.2 µM GEX1A (*Right*) (see also Supplemental Figure 2C).

After two weeks of growth on GEX1A-containing medium, plants were subjected to photorespiratory stress. Similar to the *SME1* mutation, 0.2 µM GEX1A treatment maintained higher F_v_’/F_m_’ ratios in wild-type and *cat2-2* during photorespiratory stress (Figure 2). As anticipated, GEX1A inhibited plant growth in all genotypes (Supplemental Figure 2A). In contrast to wild-type seedlings, GEX1A severely decreased F_v_’/F_m_’ in *sme1-2* seedlings already prior to stress (Supplemental Figure 2B), probably due to the combined chemical and genetic perturbation of the spliceosome. All together, these results show that *SME1* mutation and GEX1A inhibition of pre-mRNA splicing increase the resilience to photorespiratory H_2_O_2_ stress.

### *SME1* mutant plants are less sensitive to methyl viologen

After prolonged exposure to cold stress (16°C), *sme1-2* plants displayed severe developmental defects, but were phenotypically identical to wild-type plants when grown at 23°C (Capovilla et al., 2018). Growth at 20°C also resulted in smaller rosette sizes compared to wild-type plants (Huertas et al., 2019). In our growth conditions, we observed also smaller rosettes at 16°C compared to wild-type plants, while this growth phenotype was less prominent at 21°C and largely disappeared at 26°C (Figure 3A), indicating that the *SME1* mutation gradually restricts plant growth as temperature decreases. SME1 was previously reported to impact the adaptation to freezing temperatures, but did not affect plant performance under dehydration and salinity stress (Huertas et al., 2019). Given that *SME1* mutation improved resilience to H_2_O_2_-induced cell death in *cat2-2* (Figure 1), we tested the effects of the superoxide (O_2_^.-^)-producing herbicide methyl viologen (MV), osmotic stress (25 mM mannitol), and salinity (50 mM NaCl) on *sme1-2* seedling growth (Figure 3B). Based on rosette area measurements, *sme1-2* seedlings are less sensitive to MV induced growth retardation (9% decreased leaf area after 15 days) compared to wild-type plants (49% decrease; Figure 3C), but they are not affected by the salt or osmotic stress treatment. Likewise, the primary root length of *sme1-2* was less affected by MV treatment compared to wild-type (Figure 3D). A knockdown allele of the close homologue of *SME1*, AT4G30330 or *SME2* (Capovilla et al., 2018; Huertas et al., 2019), did not show phenotypical differences compare to wild-type plants under similar abiotic stress or temperature regimes (Supplemental Figure 3). In summary, *sme1-2* mutants are more resilient to MV provoked stress.

**Figure 3.**
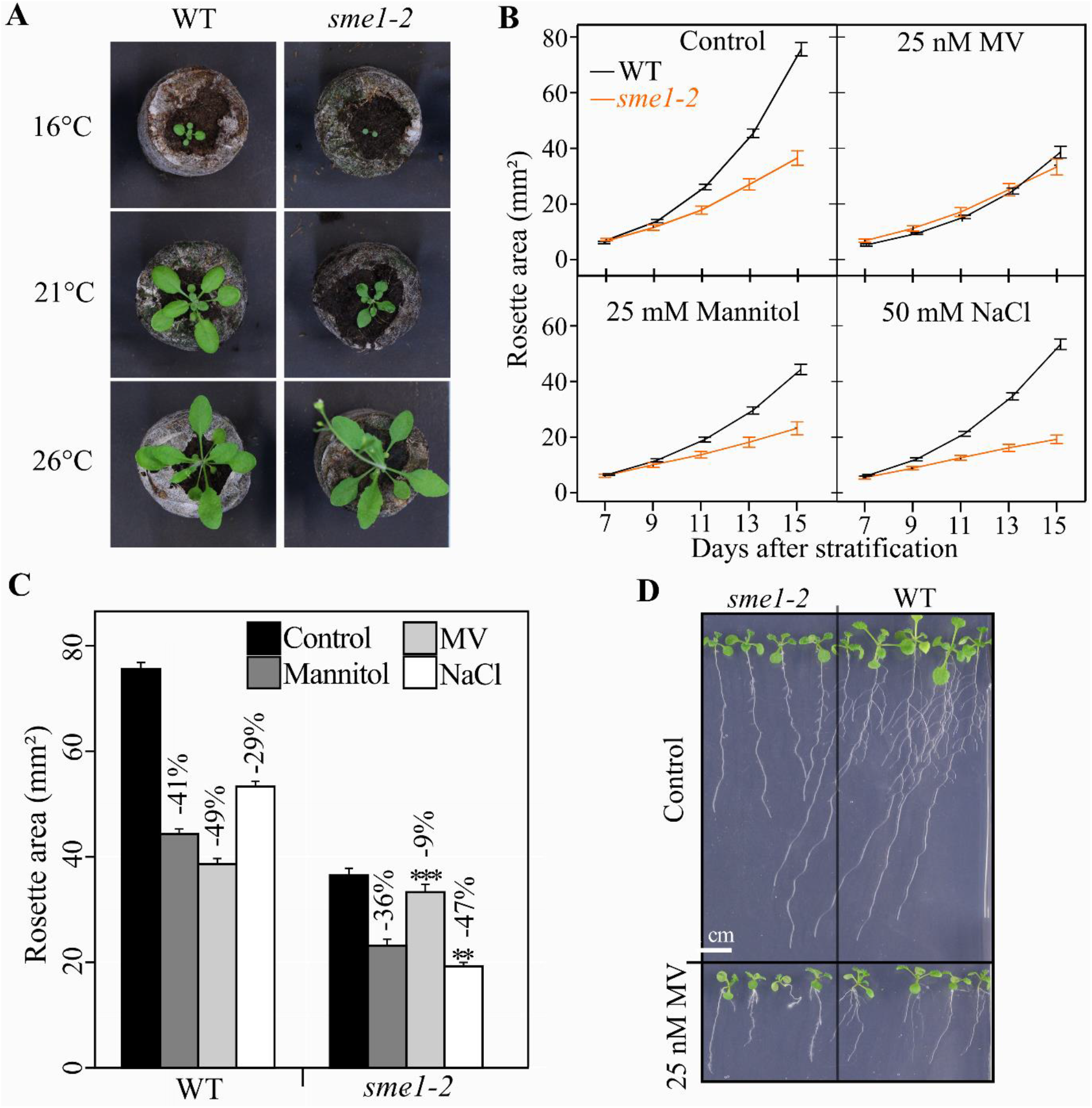
Growth-related abiotic stress assays on *SME1* mutant plants. A, Representative pictures of 3-week-old plants grown at different temperatures (16°C, 21°C, and 26°C). B, Rosette area of seedlings (n = 6, mean ± 95% confidence interval) grown on control media and media supplemented with 25 nM methyl viologen (MV), 25 mM mannitol, or 50 mM NaCl. C, Rosette area of seedlings 15 days after stratification under control and stressed conditions. Percentages indicate per genotype the rosette area reduction under stressed condition compared to control condition. Asterisks indicate significant different growth ratios of mutant versus wild-type seedlings in control and stress condition (*P*-value mixed model Wald tests, * < 0.05, * < 0.01, and *** < 0.001). D, Root growth of seedlings grown on vertical plates containing control media and media supplemented with 25 nM MV.

### *SME1* mutation triggers a constitutive molecular stress response

Next, we quantified the mRNA and protein landscapes of *sme1-2* and wild-type seedlings grown under control conditions, by mRNA-seq and liquid chromatography tandem mass spectrometry (LC-MS/MS), respectively. Compared to wild-type, *sme1-2* mutants had 2,553 induced and 887 repressed genes (Figure 4A and Supplemental Table 1; absolute fold change (FC) > 2, Q-value ≤ 0.01). Of the 4,655 proteins identified by LC-MS/MS, 314 proteins were significantly induced and 75 repressed in *sme1-2* (False discovery rate [FDR] ≤ 0.05; Figure 4B and Supplemental Table 2). Gene set enrichment analysis of both differential expressed gene (DEG) and protein lists reveals a strong enrichment of genes/proteins involved in stress and antioxidant pathways, such as glutathione metabolism (Figure 4C, Supplemental Table 3). Also secondary metabolite pathways known to stimulate protective effects under stress were differentially regulated, for instance evidenced by the strong elevated protein levels of anthocyanin biosynthetic enzymes (Supplemental Figure 4). The stress-related molecular reprogramming in *sme1-2* is further exemplified by the strong induction of pathogenesis-related (PR) genes that is a typical hallmark for the induction of a defense response (Figure 4D). We also compared the 3,440 differentially expressed genes (DEGs) identified for *sme1-2* in this study with those reported by two independently published mRNA-seq studies (Capovilla et al., 2018; Huertas et al., 2019). At a 1% FDR and a minimal 2-fold expression change (absolute log2 FC ≥ 1), Capovilla et al. reports 594 DEGs (360 induced, 234 repressed) in nine-day-old *sme1-2* seedlings grown in 16° or 23°C temperatures compared to wild-type plants. These 594 DEGs significantly overlapped with the DEGs identified here (hypergeometric *P*-value < 0.001), with 263 genes similarly down-or upregulated. In another study, Huertas et al. identified 421 DEGs in two-week-old *sme1-2* plants grown at 20°C. Of these 421 DEGs, 136 were common to this study (hypergeometric *P*-value < 0.001). Overcoming possible variations due to the mRNA-seq analysis, plant growth conditions, and other factors, a total of 382 DEGs were common to at least two of the three studies (Figure 4E). Performing GSEA with these 382 common DEGs results in an overrepresentation of several stress related gene sets (Supplemental Table 3), further consolidating that *SME1* mutation provokes a stress response in the absence of stress that probably underlies the observed resilience to H_2_O_2_-induced cell death.

**Figure 4.**
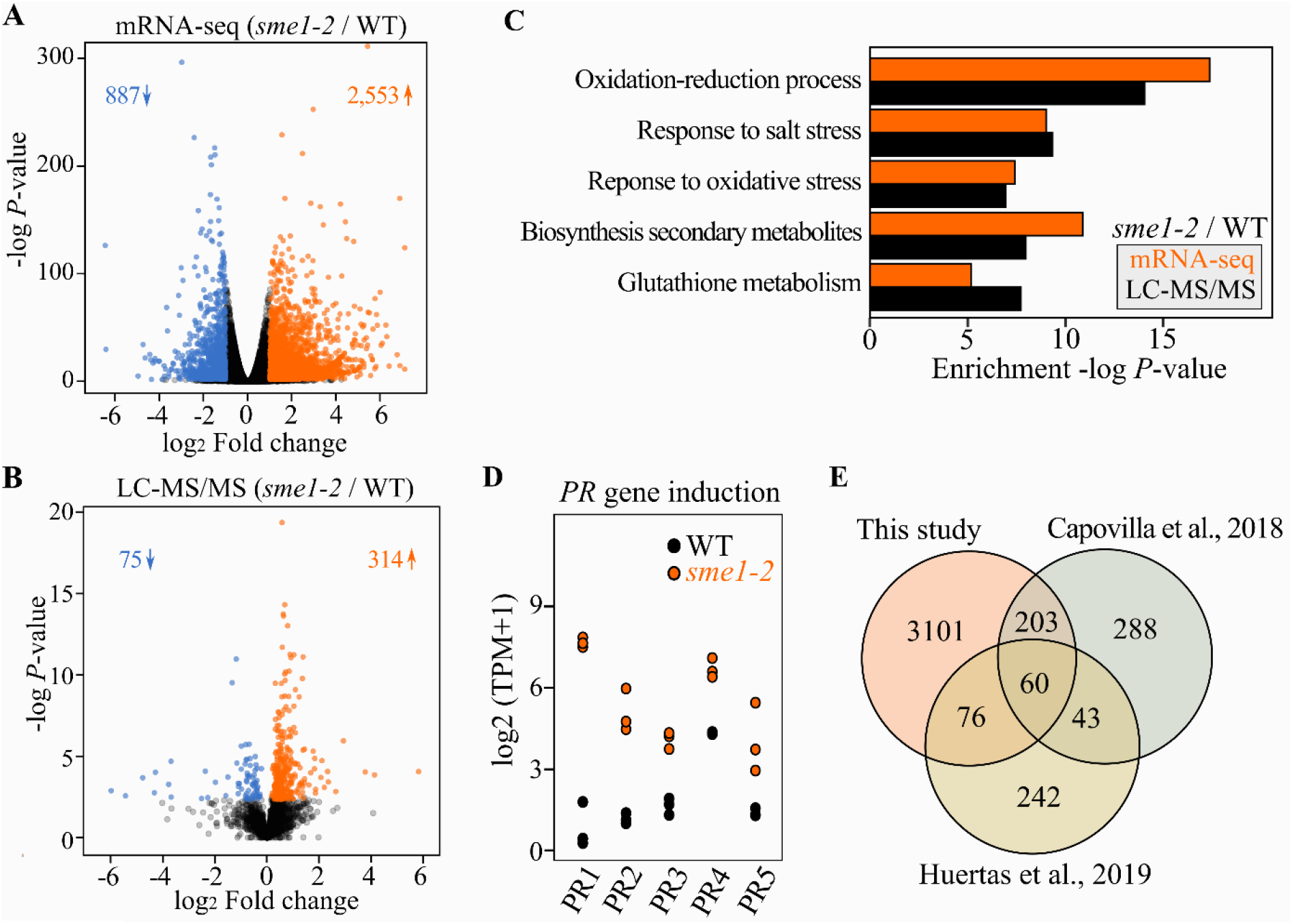
Molecular stress imprint in *sme1-2*. A, Volcano plot for differential expression in *sme1-2*. Significantly (Q-value ≤ 0.01) upregulated genes are indicated in orange (log2 FC ≥ 1) and downregulated genes in blue (log2 FC ≤ -1). B, Volcano plot for differential protein abundance. Significantly (FDR ≤ 0.05) induced and repressed proteins are indicated in orange and blue, respectively. C, Gene set enrichment analysis displaying the five most enriched gene sets in both differential gene/protein lists identified by mRNA-seq (orange) and LC-MS/MS (black), respectively (see Supplemental Table 3). D, Expression (log_2_ [transcripts per million expression+1]) of *PATHOGENESIS-RELATED* (PR) genes. E, Overlap of *sme1-2* DEGs identified in this study and those reported previously (Capovilla et al., 2018; Huertas et al., 2019).

### *SME1* mutation alters splicing patterns of metabolic enzymes and RNA-binding proteins

We assessed AS events as a consequence of *SME1* mutation by comparing mRNA-seq mapping data of *sme1-2* to wild-type. Firstly, we used the reference-based transcript assembler Scallop (Shao and Kingsford, 2017) to detect unannotated splicing junctions. After stringent filtering criteria (see Materials and Methods), an additional 16,761 Scallop-derived transcripts were added to the Araport11 annotated transcripts, which counts 48,359 protein-coding transcripts (Cheng et al., 2017). Approximately 65% of these additional Scallop-derived transcripts are probably nonfunctional because they introduce premature termination codons and display nonsense-mediated decay (NMD) features such as extended 3’ UTR lengths (Kalyna et al., 2012) (Supplemental Figure 5). Next, we determined SME1 dependent AS with SUPPA2 (Trincado et al., 2018). This method analyzes AS events such as intron retention, exon skipping, and others, based on calculations of the proportion of spliced-in (Psi) events per sample. In *sme1-2*, 785 genes with 1,114 significant AS events were uncovered when in comparison to wild-type plants (*P*-value ≤ 0.05; Supplemental Table 4), of which intron retention was the most prominent (51% of all events, Figure 5A). Next to the Psi-based AS analysis, we also analyzed differential transcript usage (DTU) in genes using the DEXSeq-stageR procedure (Anders et al., 2012; Soneson et al., 2016; Van den Berge et al., 2017), which detects shifts in the relative transcript abundance of genes. This analysis revealed 1,627 genes to show significant DTU in *sme1-2* (overall FDR ≤ 5%, Supplemental Table 5), of which 425 genes overlapped with the 785 genes with AS events identified by SUPPA2 (Figure 5B). Next, these 425 common genes, identified by both the Psi- and DTU-based analysis to undergo AS, were used for gene set enrichment analysis. In total, eight gene sets were enriched at a FDR < 5% (Figure 5C, Supplemental Table 6), of which six KEGG pathways including core metabolic pathways ‘Carbon metabolism’ (FDR 1.00^-7^), ‘Glycolysis/gluconeogenesis’ (FDR 0.0088) and ‘Biosynthesis of amino acids’ (FDR 0.013). Aside from metabolic enzymes, genes involved in response to cadmium were overrepresented, a stress intricately linked to oxidative stress (Chmielowska-Bąk et al., 2014).

**Figure 5.**
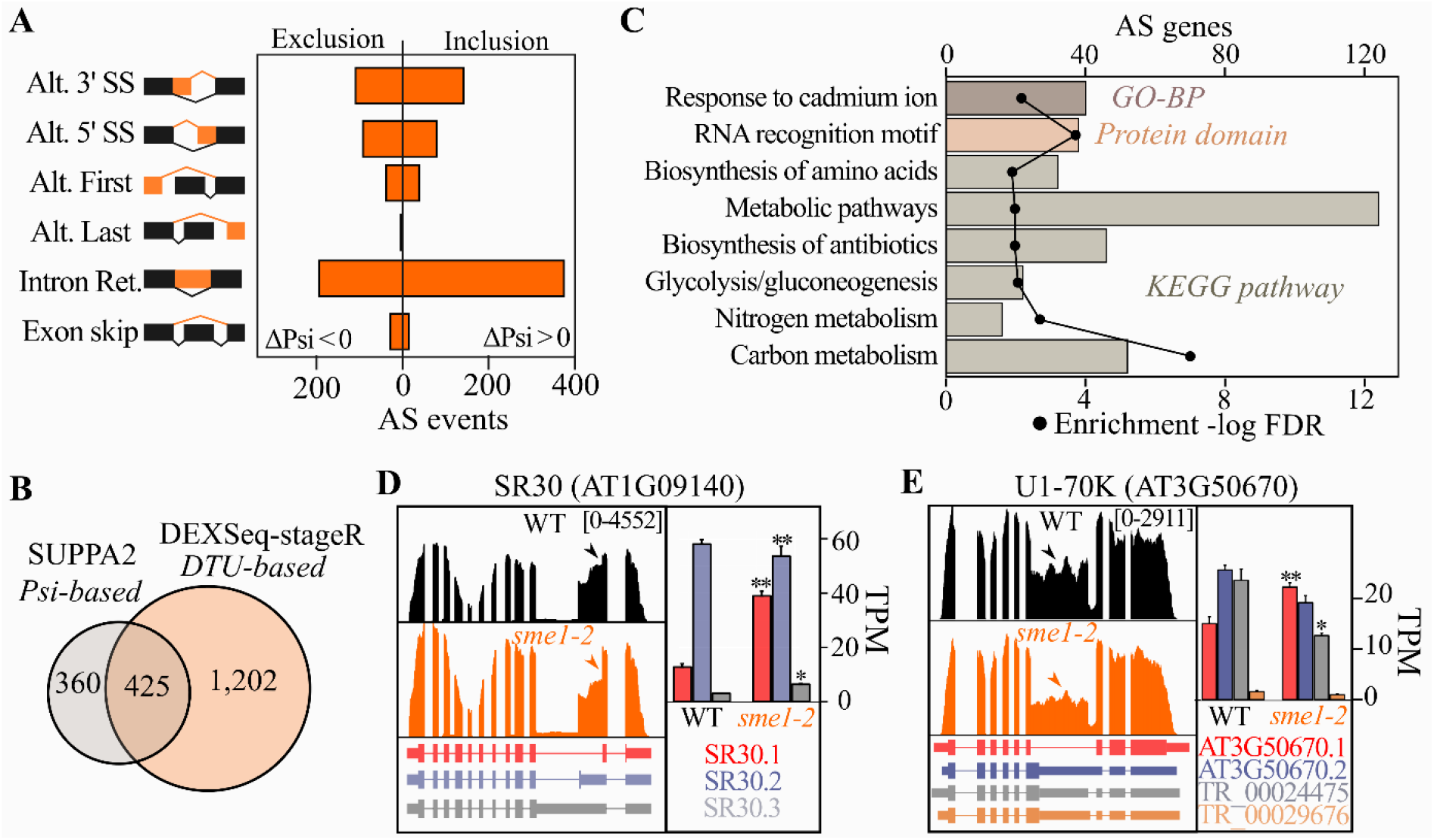
Alternative splicing analysis in *sme1-2*. A, Differential AS events (*P*-value < 0.05) identified in *sme1-2* according to splicing events detected by SUPPA2 (Trincado et al., 2018) (Supplemental Table 4). Inclusion events (ΔPsi > 0) are distinguished from exclusion (ΔPsi < 0) events. B, Overlap of identified genes with AS events detected by SUPPA2 (Trincado et al., 2018) and with significant DTU (≤ 5% overall FDR; Supplemental Table 5). C, Gene set enrichment analysis presenting gene sets enriched at a FDR ≤ 0.05 (Supplemental Table 6). Data points indicate -log_10_ FDR (lower x-axis) and bar sizes the gene count (upper x-axis). (**D-E**) Genome read coverage maps for alternatively spliced genes in wild-type (black) and *sme1-2* (orange). Corresponding transcripts were color-coded and kallisto transcript per million (TPM) quantifications represented in a bar plot (n = 3, mean ± SEM). Scallop-assembled transcripts have a ‘TR_’ prefix. Asterisks indicate stage-wise adjusted *P*-values for transcripts (* < 0.05 and ** < 0.001).

Interestingly, genes encoding proteins containing a RNA recognition motif (RRM), a prevalent eukaryotic RNA-binding domain, were also overrepresented amongst alternatively spliced genes (FDR 1.95^-4^, Figure 5C). In total, these included 19 RRM-containing proteins such as for instance the Ser/Arg-rich splicing factor SR30. Here, levels of the SR30.1 transcript (AT1G09140.1) were significantly higher in *sme1-2* compared to wild-type (Figure 5D). Interestingly, these increased SR30.1 transcript yields functional SR30 protein, unlike the SR30.2 and SR30.3 (AT1G09140.2 and AT1G09410.3) transcripts that are retained in the nucleus or are sensitive to NMD, respectively (Petrillo et al., 2014; Hartmann et al., 2018). As another example, the U1 snRNP gene U1-70K showed increased levels of the functional protein-encoding AT3G50670.1 in *sme1-2* (Figure 5E). In mammals, a similar increased splicing efficiency of the functional U1-70K is described as a compensatory mechanism upon decreased U1 snRNP protein U1C levels (Rösel-Hillgärtner et al., 2013). Other RRM containing proteins with splicing alterations included the snRNP associated proteins U2B’’, U2AF65A, RS40, RBP4C and other RNA-binding proteins (Supplemental Table 6). In summary, the *SME1* mutation results in prevalent pre-mRNA splicing alterations in genes, with overrepresentation of genes encoding metabolic enzymes and RNA-binding proteins.

### SME1 interactomics captures plant spliceosomal complexes and suggests pre-mRNA splicing related functions for uncharacterized proteins

We made fusion constructs of SME1 with a GS^green^ tag (Blomme et al., 2017), and produced transgenic plants to assess the SME1 protein subcellular localization. Confocal microscopy indicated that SME1 has a predominant nuclear localization in Arabidopsis (Figure 6A and 6B).

**Figure 6.**
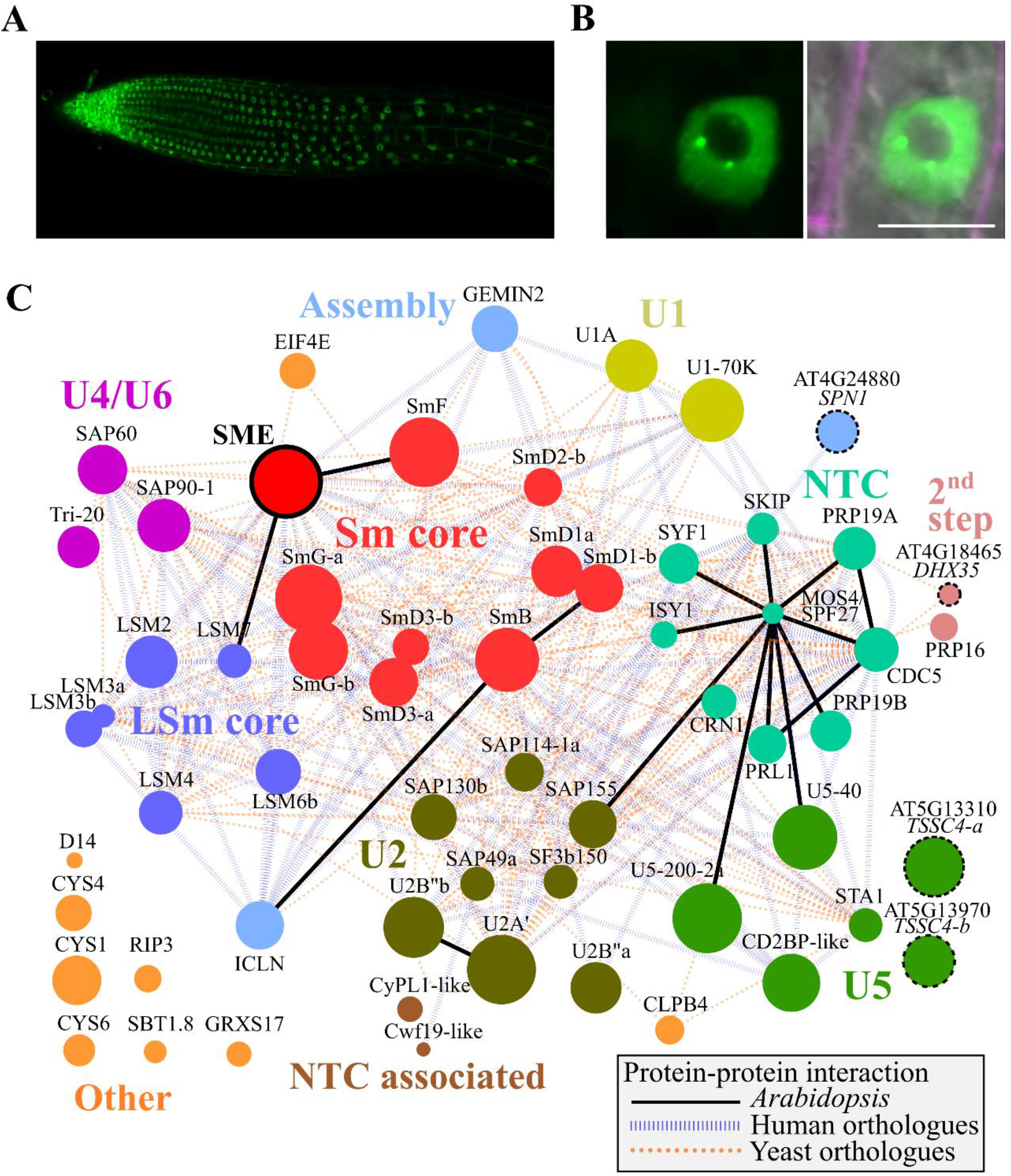
Nuclear localization and SME1 interactome. Confocal microscopy of N-terminal SME1-GS^green^ construct in whole root A, and representative root tip cell B,. For the root tip cell an overlay of GFP signal, propidium iodide (PI) counterstain, and brightfield is displayed. Scale bar: 10 µm. C, SME1 protein interaction network. All 60 proteins identified by TAP-MS (Supplemental Table 7) are displayed as nodes sized according their protein LFQ intensity and colored by their respective spliceosome complex (Koncz et al., 2012). In addition, GEMIN2, ICLN and SPN1 were categorized as spliceosome assembly proteins and four uncharacterized plant proteins (encoded by AT4G18465, AT4G24880, AT5G13310, and AT5G13310 – dotted borders) show homology to known pre-mRNA splicing proteins (see text). Edges reflect protein interactions previously reported between *Arabidopsis* proteins (solid black lines), their human orthologs (blue vertical slashes), or yeast orthologs (orange dashes) in the IntAct database (Hermjakob et al., 2004).

While decreased signal is apparent in the nucleolus, some nuclear speckles are highlighting (Figure 6B), hence indicating a spatial correlation between SME1 and matured spliceosomal snRNPs that facilitate pre-mRNA splicing (Matera and Wang, 2014). Next, we assessed the SME1 interactome by tandem affinity purification-mass spectrometry (TAP-MS) in transgenic Arabidopsis cells expressing SME1 fused to a GS^rhino^ TAP tag (Van Leene et al., 2015). In total, 60 proteins were identified (Supplemental Table 7) in a complex trapped by the SME1 fusion protein. We generated the SME1 interactome network, also including previously reported protein-protein interactions (Figure 6C – network edges) between the 60 SME1 protein interactors (network nodes) in Arabidopsis, as well those reported between their respective human and mouse orthologs. The SME1 interactome entails the six other Sm proteins that form the heptameric Sm protein complex. In addition to the Sm proteins, also five out of the eight LSm proteins were detected, together with other associated proteins of the U1 (2 proteins), U2 (7 proteins), U4/U6 (3 proteins), and U5 snRNP (4 proteins) complexes (Supplemental Table 7) and eight proteins of the NineTeen Complex (NTC), a complex involved in the remodeling of spliceosomes during pre-mRNA splicing (Matera and Wang, 2014). In addition, two other proteins that have been implicated in assembly of snRNP complexes are also part of the SME1 interactome. AT1G54380 encodes GEMIN2 that is part of the survival motor neuron (SMN) complex governing snRNP core assembly (Zhang et al., 2011; Schlaen et al., 2015). Arabidopsis *GEMIN2* mutants display pre-mRNA splicing defects that impact the circadian rhythm and cold acclimation (Schlaen et al., 2015). In addition, AT5G62290 encodes ICLN that is an ortholog of human pICln (or CLNS1A), an assembly chaperone that recruits Sm proteins to the PRMT5 methylation complex prior to delivery to the SMN complex (Chari et al., 2008) and shown to be associated with PRMT5 and SmD3b in Arabidopsis (Huang et al., 2016).

Lastly, we identified four SME1 protein interactors that are orthologs to human splicing related genes for which, to our knowledge, no functional characterization is reported in plants. Firstly, AT4G24880 is an ortholog of human Snurportin-1 (SPN1) that catalyzes the nuclear import of snRNPs (Huber et al., 1998) (Supplemental Figure 6A). In addition, AT4G18465 is a DEAH box RNA helicase with best homology to human DHX35 (Supplemental Figure 6B, BLASTP E-value < 1e^-100^), a helicase which associates with the human spliceosome and likely involved in second step chemistry during splicing (Ilagan et al., 2013). Lastly, the paralogs AT5G13310 and AT5G13970, encoded from a duplicated gene pair, were both identified as SME1 interactors. Although showing no apparent sequence similarity to mammalian splicing factors, both proteins contain a ‘Tumor suppressing sub-chromosomal transferable candidate 4’ (TSSC4) domain, derived from the human TSSC4 protein that was recently identified as a new chaperone that acts in U5 snRNP de novo biogenesis as well as post-splicing recycling (Klimešová et al., 2021). I Similar as in a previous conservation analysis, multiple sequence alignment indicates that the structured Hom3, contributing to interacting with the U5 snRNP complex, shows the strongest conservation in plant TSSC4 proteins (Supplemental Figure 7A). In addition, like in the human TSSC4, ∼70% of the protein structure of both TSSC4 proteins in Arabidopsis is predicted to be disordered (Supplemental Figure 7B). On basis of these sequence characteristics and its identification in the SME1 interactome, we designated this gene pair as TSSC4-a and TSSC4-b (Supplemental Table 7). Of the remaining nine protein interactors, also the eukaryotic translation initiation factor EIF4E was previously shown to physically interact with spliceosomal complexes and influence splicing in human cell lines (Ghram et al., 2022). Surprisingly, three cystatin protease inhibitors (CYS1, CYS4 and CYS6) and the protease SUBTILASE1.8 were identified as interactors. Another protease, METACASPASE1, was recently discovered to associate and stabilize LSm4 in Arabidopsis (Wang et al., 2021) and tobacco cystatins have been observed in the nucleus (Zhao et al., 2014). As such, it is plausible that additional proteases and/or their inhibitors interact with plant spliceosomes. Altogether, these data clearly illustrate that snRNP complexes and associated proteins were succesfully captured by the TAP-MS analysis and enabled the identification of proteins that were previously not known to be involved in pre-mRNA splicing in Arabidopsis.

### Sm ring assembly mutants show altered sensitivity to oxidative stress

Mutation of *SME1* and chemical inhibition of pre-mRNA splicing alleviated in *cat2-2* cell death alleviation, suggesting that other mutations that affect spliceosome function could yield resistance to oxidative stress. Given that ICLN acts in snRNP assembly (Chari et al., 2008), we tested whether its loss-of-function (*icln-2*, SALK_050231C; Huang et al., 2016) resulted in similar phenotypes as *SME1* mutation. Firstly, we monitored U snRNA levels. Similar to *gemin2* and *sme1-2* mutants (Huertas et al., 2019; Schlaen et al., 2015), drastic U2 snRNA and mild U1 and U4 snRNA decreases were observed in *icln-2* (Figure 7A). As it is known that impairment of snRNP complexes results in destabilization of snRNA, this provides further argumentation for ICLN as a functional snRNP assembly protein ortholog in plants. Moreover, *icln-2* accumulated increased levels of SME1, hinting at a possible compensatory effect (Figure 7A). Besides decreased growth at lower temperatures, *icln-2* displayed some mild developmental defects at 21°C with shortened petioles and a more compact leaf shape (Figure 7B). In addition, when transferring one-week-old *icln-2* seedlings from 21°C to 10°C, these were less retarded in growth than *sme1* seedlings (Figure 7C). *icln-2* seedlings were also subjected to abiotic stresses. Like *sme1-2, icln-2* leaf area was significantly less impacted by MV stress (−28% versus -51% in wild-type), whereas salt or mannitol stress caused decreased growth rates similar to wild-type (Figure 7D, Supplemental Figure 8). Taken together, albeit less pronounced, *icln-2* shows many phenotypical similarities to *sme1-2*, suggesting impaired snRNP assembly and SME1 mutation to result in developmental defects at lower temperature and increased resilience to oxidative stress.

**Figure 7.**
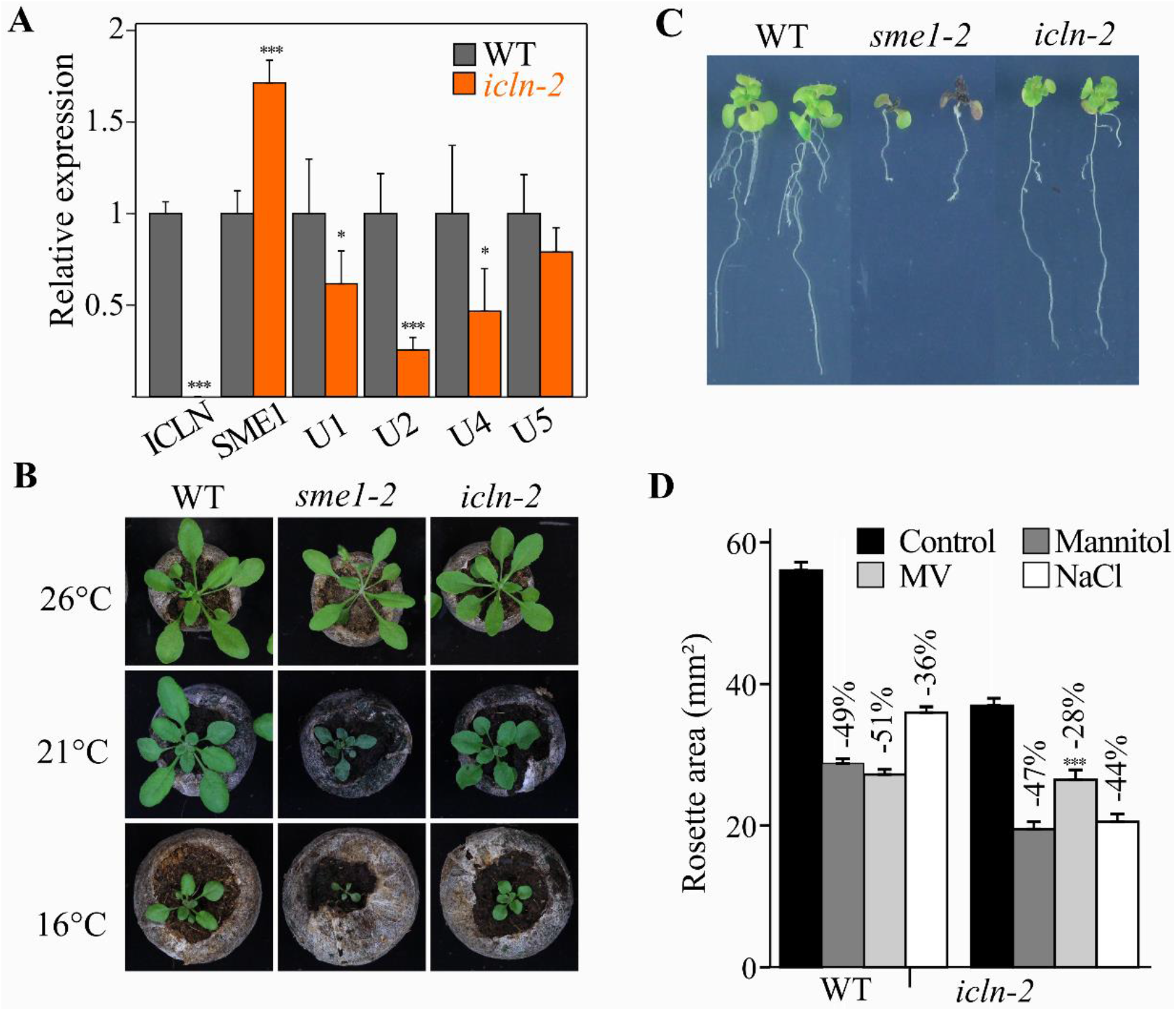
Mutation of ICLN results in similar phenotype as SME1 mutation. A, Relative transcript levels of *ICLN, SME1*, and U1, U2, U4, U5 snRNAs measured by qPCR in wild-type and *icln-2* seedlings (n = 3, mean ± SEM). B, Representative pictures of 3-week-old plants grown at 26°C, 21°C, or 16°C (top to bottom). C, One-week-old seedlings grown under control conditions (21°C) for one week on vertical plates and shifted to 10°C. Pictures were taken 4 weeks after transfer. D, Rosette area of seedlings 15 days after stratification grown on control media and media supplemented with 25 nM methyl viologen (MV), 25 mM mannitol, or 50 mM NaCl. Percentages indicate per genotype the rosette area reduction under stressed condition compared to control condition. Asterisks indicate significant different growth ratios of mutant versus wild-type seedlings in control and stress condition (*P*-value mixed model Wald tests, * < 0.05, ** < 0.01, and *** < 0.001).

Lastly, we analyzed leaf area under abiotic stresses of T-DNA insertion lines of the four SME1 interactors SPN1, DHX35, TSSC4a and TSSC4b that showed homology to mammalian pre-mRNA splicing proteins (Supplemental Table 7). Altered stress tolerance could further hint towards a functional pre-mRNA splicing role of these proteins in Arabidopsis. A SPN1 knockdown allele, which maintained ∼37% expression (Supplemental Figure 9), showed a significant decreased leaf area compared to wild-type under MV stress (−61% versus -46% in wild-type, *P*-value 2.4E-04) after 17 days, but not salt or mannitol stress (Supplemental Figure 8). In contrast, loss-of-function alleles for DHX35, TSSC4a and TSSC4b (with 5-8% remaining expression, Supplemental Figure 9) showed no significant leaf area different under normal or abiotic stress conditions (Supplemental Figure 8). In addition, no significant U snRNA levels were recorded, except for decreased U2 snRNA levels for the DHX35 mutant (Supplemental Figure 9B).

## DISCUSSION

Alternative splicing is an important posttranscriptional regulation mechanism in diverse plant processes, including abiotic stress responses (Laloum et al., 2018), plant immunity (Rigo et al., 2019), circadian rhythm (Nolte and Staiger, 2015), and plant development (Szakonyi and Duque, 2018). Accordingly, mutations of core snRNP proteins, e.g. Sm or LSm proteins, or proteins facilitating snRNP assembly, result in pleiotropic developmental phenotypes, and altered stress tolerance and circadian rhythms (Swaraz et al., 2011; Schlaen et al., 2015; Huang et al., 2016; Carrasco-López et al., 2017). Two recent publications described growth and developmental defects of *sme1-2* mutants and the role of SME1 in pre-mRNA splicing and cold acclimation (Capovilla et al., 2018; Huertas et al., 2019). Thus far, plant splicing mutants were only anecdotally correlated with altered oxidative stress responses. A mutation of the U1 specific snRNP protein U1A showed an increased ROS accumulation and hypersensitivity to salt stress and growth on MV decreased primary root length when compared to the wild-type, whereas overexpression lead to an, albeit moderate, increased resilience against salt stress (Gu et al., 2018). Here, we identified a nonsense mutation in *SME1* to alleviate H_2_O_2_-induced cell death in *cat2-2*. The C-terminal region of SME1 was already shown to be essential for its function (Capovilla et al., 2018), in line with the 67 AA C-terminal truncation presented here by the EMS-induced mutation. Next to *SME1* mutation, application of the spliceosome inhibitor GEX1A attenuated the *cat2-2* cell death phenotype, suggesting pre-mRNA splicing inhibition as major causal factor in *SME1* mutants. Further phenotypical screening revealed *SME1* mutation to render plant seedlings leaf and root growth more tolerant to MV, unlike salt or mannitol stress. Together with the earlier observed decreased sensitivity to photorespiratory-driven H_2_O_2_, this suggests *SME1* mutation to exhibit an improved resilience to increased superoxide (O_2_^.-^) levels.

This improved resilience is in agreement with the differential gene and protein expression recorded by mRNA-seq and proteomics, where a constitutive stress response is apparent in the absence of stress (Figure 4C and 4D, Supplemental Table 3). These difference could be attributable to numerous reasons such as plant tissue, condition, sample variation or used algorithms. For instance, our employed statistical software for differential testing, i.e. kallisto-sleuth (Bray et al., 2016; Pimentel et al., 2017), cannot process single replicate data as analyzed in the mRNA-seq study of Huertas et al. Other differentially regulated processes included glutathione metabolism and secondary metabolism induced enzymes involved in the production of anthocyanins that protect against oxidative stress and typically give rise to a darkened leaf color clearly visible under the low temperature shift of *sme1* (Figure 7C). It should be noted that the amount of DEGs in this study far exceeds those reported before, i.e. over 3,000 DEGs in this study versus approximately 400 and 600 DEGs at similar FDR and log2 FC thresholds (Capovilla et al., 2018; Huertas et al., 2019).

As described in Arabidopsis and human cell lines (Chen et al., 2019; Huertas et al., 2019), intron retention was identified in this study as the major AS splicing event after SME1 depletion (Figure 5A). In this study, SME1 mutation dependent AS was overrepresented in genes involved in metabolic pathways and encoding RNA-binding proteins (Figure 5C). Cross-or auto-regulation via AS of pre-mRNA splicing genes has been described before under temperature changes (Verhage et al., 2017). Also in animals, splicing factors, such as SR proteins, are known to exert auto-regulatory feedback by unproductive splicing and hence regulate the fate of their own mRNAs (Müller-McNicoll et al., 2019). Here, we also observed compensatory splicing of spliceosome components, such as U1-70K, as previously described in the Arabidopsis Sm core assembly mutant *gemin2* (Schlaen et al., 2015). Nevertheless, it remains challenging to pinpoint which AS events precisely contribute (partly) to the observed stress reprogramming and/or developmental phenotypes. Notably, AS in the loss-of-function Arabidopsis mutant U1A affected the redox-related genes *ACONITASE1* and *COPPER/ZINC SUPEROXIDE DISMUTASE 1* which resulted in excess ROS accumulation and increased sensitivity to salt stress (Gu et al., 2018). While no similar intron retention was observed for *ACONITASE1* (Supplemental Figure 10A), we observed retention of intron 4 and 7 in *sme1-2* (Supplemental Figure 10B), while intron 6 was retained in the *atu1a* mutant.

Amongst the protein interactors isolated with SME1 as bait in a pull-down experiment, we identified the snRNP assembly proteins GEMIN2 and ICLN (Huang et al., 2016; Schlaen et al., 2015). ICLN is together with PRMT5 part of the methylosome complex that methylates Arg residues of SmB, SmD1, and SmD3 and delivers Sm protein complexes to the survival motor neuron (SMN) complex (Matera and Wang, 2014). Although it was already shown that ICLN interacts with SmD3-b and PRMT5 *in planta* (Huang et al., 2016), we further support its role in snRNP assembly in this study as *icln-2* mutants display decreased U1, U2, and U4 snRNA levels (Figure 7A). Reduced U snRNA levels were observed earlier in ICLN depleted *Saccharomyces pombe* (Barbarossa et al., 2014). However, Arabidopsis ICLN could not complement the fission yeast mutant (Huang et al., 2016), unlike the human pICLN protein (Barbarossa et al., 2014) – indicating functional differences of Arabidopsis ICLN. In this study, we further established growth of *icln-2* to be less sensitive to MV stress (Figure 7D), further suggesting that suboptimal snRNP assembly – i.e. also improper SME protein levels, underlie the observed resilience to oxidative stress resilience. Methylosome components SmD3b, ICLN2 and PRMT5 are negatively regulating plant immunity as *icln-2, prmt5*, and *smd3b* mutants show an increased resistance to oomycete pathogens (Huang et al., 2016). In addition, the PR gene induction in *sme1-2* was also observed earlier in *icln-2* and *smd3b* mutants (Huang et al., 2016). Subsequently in snRNP assembly, the SMN member GEMIN2 binds directly to Sm proteins to facilitate further snRNP assembly (Matera and Wang, 2014). Aside from reduced U snRNA levels, *gemin2* mutants were affected in circadian rhythm linked to mis-splicing of core clock genes, such as, for instance, retention of intron 4 of *CIRCADIAN CLOCK ASSOCIATED 1* (*CCA1*), which was also observed in *sme1-2* (Supplemental Figure 10C). This intron retention in *CCA1* gives rise to the truncated CCA1β splice form that interferes with the formation of functional CCA1 complexes (Seo et al., 2012). Aside from *CCA1*, other clock genes were overrepresented as alternatively spliced genes in *sme1-2* (Supplemental Table 6). Notably, the tolerance to a foliar 5 µM MV spray was enhanced in *CCA1*-overexpressing plants, whereas *cca1* and other clock gene mutants were hypersensitive to MV (Lai et al., 2012). Hence, disrupted circadian rhythm due to AS of clock genes might (partly) be associated with the observed resilience to oxidative stress in *sme1*.

Given that 47 out of 60 SME1 protein interactors are splicing-related proteins (Supplemental Table 7), we hypothesized that the other SME1 protein interactors might have uncharacterized roles in pre-mRNA splicing. One of these interactors is homologous to human SPN1 (Supplemental Figure 6A), which facilitates the nuclear import of snRNPs by binding to the 2,2,7-trimethylguanosine (m3G)-cap of U snRNAs (Huber et al., 1998) and directly interacts with human SmB and SmD3 (Kühn-Hölsken et al., 2010). Whereas no developmental phenotypes or altered U snRNA levels were observed for the *spn1* mutant in this study, growth of *spn1* seedlings is significantly decreased on MV-supplemented media compared to wild-type (Supplemental Figure 8). Lastly, we identified an uncharacterized Arabidopsis gene pair encoding proteins containing TSSC4 domain, which was recently linked to tri-snRNP maturation as U5 snRNP component (Klimešová et al., 2021). Although the low overall sequence conservation between both proteins, explained by the multiple disordered regions, the Hom3 domain involved in U5 snRNP binding showed the highest degree of conservation in plants (Supplemental Figure 7). No phenotypical differences to wild-type were uncovered, possibly due to a functional redundancy between both genes. To tackle this a double mutant is being generated, but which is challenged by the proximity of both genes on the chromosome. Although the observed analogy with human proteins and (indirect) interaction with SME1 in Arabidopsis, further follow-up studies are required to verify their role in pre-mRNA splicing.

Regulation of snRNP assembly and pre-mRNA splicing activity are correlated with adaptive responses. For instance, the expression of *GEMIN2, SME1* and *LSm* genes is transcriptionally upregulated during cold stress (Capovilla et al., 2018; Huertas et al., 2019; Schlaen et al., 2015; Carrasco-López et al., 2017; Perea-Resa et al., 2016). Moreover, the increased U1 and U6 snRNA levels under cold stress were attributed either to enhanced accumulation of snRNP complexes or to a compensatory effect in response to reduced snRNP functionality (Carrasco-López et al., 2017; Schlaen et al., 2015). Besides enhanced expression, SME1 preferentially localizes in the nucleus under lower temperatures (Huertas et al., 2019) and also LSM8 GFP signals are enhanced under cold (Carrasco-López et al., 2017). Posttranslational modifications (PTMs) are yet another regulation layer that could influence spliceosome stability or activity. For instance, the protease METACASPASE1 associates with LSm4 and affects AS of pre-mRNAs (Wang et al., 2021), thus indicating proteolysis in proximity of spliceosome complexes as putative regulatory mechanism. PTMs are of particular relevance in the context of H_2_O_2_ signaling are oxidative PTMs on Cys residues (Akter et al., 2015). For instance, in human, snRNP assembly is known to be inactivated by oxidative stress due to intermolecular disulfide formation of the SMN protein (Wan et al., 2008), which however lacks a true ortholog in Arabidopsis (Schlaen et al., 2015). In Arabidopsis, cysteine redox signaling has previously been identified at the snRNP assembly methylosome complex that methylates Sm proteins and delivers them to the SMN complex. More specifically, S-nitrosylation of PRMT5 Cys216 increased PRMT5 methylation activity and facilitated stress dependent splicing and tolerance to salt (Hu et al., 2017). In a recent chemoproteomic study, the same PRMT5 Cys216 showed a 2.6-fold increased oxidation upon H_2_O_2_ treatment in cell cultures (Huang et al., 2019). Redox-sensitive Cys residues were also identified previously for two Sm proteins, SmB and SmD2. In the case of SmD2, a 2.4-fold increased oxidation was apparent for Cys41 and Cys58 under H_2_O_2_ stress (Huang et al., 2019). Moreover, both Cys were captured *in vivo* by a redox-active Yap1 construct in Arabidopsis (Wei et al., 2020) and were similarly oxidized in the human ortholog SNRPD2 (Yang et al., 2014; Gupta et al., 2017). Hence, oxidative PTMs could potentially alter snRNP assembly and/or functionality in plants, resulting in pre-mRNA splicing inhibition and defensive mechanisms in return.

## MATERIALS AND METHODS

Details regarding proteome extraction, LC-MS/MS, and protein database searches are included in the Supplemental Experimental Procedures.

### Plant material

T-DNA insertion lines SALK_089521 (*sme1-2*), SALK_050231C (*icln-2*), and SALK_119615 (*spn1*), SALK_139760 (*sme2*), SALK_007841C (*dhx35*), SALK_047869C (*tssc4a*), and SALK_118948C (tssc4b) were obtained from the Nottingham Arabidopsis Stock Center. T-DNA insertion position was confirmed with gene-specific and T-DNA-specific primers (Supplemental Table 8) with Phire Plant Direct PCR Master Mix (Thermo Scientific) according to the manufacturer’s guidelines.

### Plant growth conditions and stress treatments

The suppressor mutation screen and photorespiratory stress treatment were done as described (Kerchev et al., 2016; Waszczak et al., 2016). Seedlings were grown on half-strength Murashige and Skoog (MS) medium under control conditions. The photorespiratory stress treatment, also termed restricted gas exchange-continuous light (RGCL), subjects plants to a continuous moderate light regime while blocking air exchange between the petri dish and the environment. Changes in F_v_’/F_m_’ signals were quantified by an Imaging PAMM-series (MAXI version) chlorophyll fluorometer and the ImagingWin software application (Heinz Walz). For the plate based oxidative stress assay, seeds were first gas sterilized, stratified for 3 days and germinated on growth medium containing half-strength MS salts, 1% (w/v) sucrose, 100 mg L^−1^ myo-inositol, 500 mg L^−1^ MES, 0.5 mg L^−1^ nicotinic acid, 0.5 mg L^−1^ pyridoxine, 1 mg L^−1^ thiamine, and 0.7% (w/v) plant tissue culture agar (Duchefa), either supplemented or not with 25 nM MV (Acros Organics) at 21°C under long-day conditions (16 h light [60-80 μmol.m^−2^.s^−1^]/8 h dark). For the cold stress experiment, seedlings were grown under a continuous light regime in a controlled environment thermostat cabinet (Lovibond) for five days at 21°C, whereafter the temperature was decreased to 10°C for 4 weeks. In soil, plants were grown at 16°C, 21°C, or 26°C, plants in controlled environment thermostat cabinets (Lovibond) in long-day regime (16 h light [90-110 μmol.m^-2^.s^-1^]/8 h dark). For growth at low and moderate light intensities (100 and 200 μmol.m^−2^.s^−1^, respectively), plants were grown in a controlled-environment growth chamber (Weiss Technik; 16 h light/8 h dark, 21°C, 60-65% humidity).

### Rosette area measurements and statistics

For rosette area quantification, plants were photographed and the images were quantified with ImageJ (version 1.45), statistical analysis was done with SAS (version 9.4). Briefly, the log-transformed rosette area was longitudinally analyzed. The Toeplitz covariance matrix was used to calculate the residual covariance. The fixed effects contained the main effects of the tested line, treatment and time, and all possible interactions. Kenward-Roger approximation was applied to determine the denominator degrees of freedom for the tests of fixed effects. Finally, user-defined contrasts were estimated with Wald tests. The contrasts of interest were the rosette area differences upon each stress compared to the control condition between each line and the control line at each day. *P*-values were appropriately adjusted with the MaxT method as implemented in SAS. Assumptions were verified through residual analysis. Model fitting and hypothesis testing were done with the mixed and the plm procedure, respectively.

### Causative mutation in *40*.*2*

To identify the causative mutation in *40*.*2*, we used the SHOREmap backcross pipeline as described (Hartwig et al., 2012; Kerchev et al., 2016; Waszczak et al., 2016). We backcrossed *40*.*2* with *cat2-2* lines and subjected the obtained F2 mapping population to the photorespiratory stress assay. Approximately 14% of the plants reversed the *cat2-2* cell death phenotype. A low segregation ratio could be due to additional EMS mutations that impact on plant growth or limited penetrance of the revertant phenotype (Kerchev et al., 2016). Afterward, 60 surviving plants were analyzed by next-generation sequencing, leading to an average 39.3-fold genome coverage allowing identification of single nucleotide polymorphisms (SNPs) associated with the cell death alleviation. A mutation hotspot was apparent in chromosome 2 and SNPs in four candidate genes showed a frequency of 100% in reversion plants (Supplemental Figure 1), among which only *SME1* contained a nonsense mutation theoretically leading to a truncated protein.

### Tandem affinity purification and data analysis

Wild-type *Arabidopsis thaliana* (L.) Heynh. (accession Columbia-0 [Col-0]) cell suspension cultures grown in a 16-h photoperiod were transformed with N-terminal GS^rhino^-tagged (Van Leene et al., 2015) SME1 constructs by *Agrobacterium tumefaciens* cocultivation. In total, in each of the two individual TAP experiments, the SME1-GS^rhino^ recombinant protein and its interacting partners were purified. Protein extract preparation, TAP, proteolysis, and mass spectrometry analysis were performed as described (Van Leene et al., 2015). Raw mass spectrometry data files were processed by MaxQuant (version 1.5.5.1) (Cox and Mann, 2008). Data were searched against the Arabidopsis proteome concatenated with protein contaminants of TAP and general proteomic contaminants (Van Leene et al., 2015). Carbamidomethyl was set as a fixed modification for Cys, whereas Met oxidation, protein N-terminal acetylation, and methylation of Asp and Glu were variable modifications. Matching between the two replicates was enabled, with an alignment window of 20 min and 0.7 matching time window. The resulting protein identification table (“proteinGroups.txt”) was filtered for contaminant and decoy protein hits. In addition, aspecific background proteins frequently identified in previous Arabidopsis TAP experiments were omitted (Van Leene et al., 2015). Protein group identification required at least one unique peptide, resulting in a list of 60 protein groups (Supplemental Table 7). The mass spectrometry proteomics data have been deposited to the ProteomeXchange Consortium (http://proteomecentral.proteomexchange.org) via the PRIDE partner repository (Vízcaino et al., 2013) with the dataset identifier PXD016607, accessible to reviewers using the user name ‘’ and password ‘9YumqPwo’.

### Real-time quantitative PCR

Total RNA was isolated with the TriZol reagent (Ambion) and the RNeasy Plant Mini Kit (Qiagen) with on-column gDNA digest with RQ1 DNase (Promega). One µg of total RNA was used for first-strand cDNA synthesis with the iScript cDNA Synthesis Kit (Bio-Rad) and diluted 8 times afterwards. Finally, 10% of the total reaction volume was comprised by the first-strand cDNA template. Real-time PCR was done with the SYBR Green I Master kit (Roche), according to the manufacturer’s instructions. Primer sequences are provided in Supplemental Table 9. Gene expression was quantified with the qBASEPlus software (Biogazelle) and the PROTEIN PHOSPHATASE 2A SUBUNIT A3 (AT1G13320) and UBIQUITIN-CONJUGATING ENZYME 10 (AT5G53300) as reference genes. Unless otherwise stated, all experiments were done in 3 biological replicates, each consisting of at least 10 plants.

### Confocal microscopy

35S:SME1-overexpressing constructs were tagged N-terminally and C-terminally with the GS^green^ tag (Blomme et al., 2017). More precisely, the coding sequence of SME1 was amplified by PCR from reverse-transcribed RNA extracted from Arabidopsis Col-0 seedlings. using gene-specific primers containing minimal AttB-sites. The obtained PCR fragments were recombined into pDONR221 with the Gateway system (Invitrogen) and subsequently into the pB7m34GW (C-terminal tag, 10524.7.6) or pB7m24GW2 (N-terminal tag, 10523.8.6) destination vectors. To generate stable transgenic lines, Arabidopsis plants were transformed by the floral dip method and propagated until homozygous T3 lines were generated. Overexpressing plants were investigated with a LSM 710 confocal microscope (Zeiss). Before the GFP expression profile check, plants were incubated for a few minutes in 1% propidium iodide for membrane staining.

### High-throughput sequencing

#### RNA extraction and RNA sequencing

RNA was isolated from shoot tissue of 3-week-old seedlings with the TRIzol Reagent (Invitrogen). Sequencing libraries were constructed with the TruSeq Stranded LT mRNA Library Preparation Kit (Illumina). The NextSeq 500 High sequencer (Illumina) was used to generate 150-bp paired-end reads at the VIB Nucleomics Core facility (http://nucleomics.be/). Demultiplexing and FASTQ data were generated on BaseSpace (Illumina’s cloud-base resource). Raw sequencing data are deposited on gene expression omnibus (GEO, https://www.ncbi.nlm.nih.gov/geo/) with the identifier GSE133182, accessible to the reviewers using the secure token ‘mpmpoykkzxqffeh’.

#### Genome alignment and transcript assembly

RNA-seq reads were mapped to the Arabidopsis genome by means of the Spliced Transcripts Alignment to a Reference (STAR) software (version 2.5.2b) (Dobin et al., 2013) using the Araport11 annotation as reference (Cheng et al., 2017). Reads were required to match uniquely to single genome locations with a 4% mismatched bp per read allowance (−-outFilterMismatchNoverLmax 0.04). STAR was run in the 2-pass mode to achieve a more sensitive discovery of splice junctions. Per sample, between 41 to 47 million 2 × 150 bp were uniquely mapped (∼ 99% input reads) to the Arabidopsis genome (Supplemental Table 10). To assemble *de novo* transcripts, read alignments (.BAM) files were used for transcript discovery with Scallop (version 1.3.3b) (Shao and Kingsford, 2017) with the option “-min_transcript_coverage 10”. In addition, in-house Perl scripts were used to filter transcripts with noncanonical splice junctions not supported by at least 20 unique reads in at least 3 samples and transcripts were required to uniquely contain exon-intron junctions of a single Araport11 gene locus to avoid merging exons of different gene loci. Genome visualizations of read coverage and transcripts structures were generated with the Integrative Genomics Viewer (version 2.4.6) (Robinson et al., 2011).

#### Transcript quantification and differential analysis

For transcript and gene quantification, we used the pseudo-alignment tool Kallisto (version 0.43.0) (Bray et al., 2016). Initially, a Kallisto index was created with Araport11 and the 16,761 novel assembled transcript sequences. Reads were quantified with 100 bootstraps and bias correction (“-b 100 --bias”). For differential analysis, the companion tool sleuth (version 2.90.0) (Pimentel et al., 2017) was used and a basic filter of at least 10 read counts for 47% of the samples. A separate model was fitted for each pairwise comparison of interest and differential expression was detected with a Wald test. Differentially expressed genes (DEGs) and transcripts were defined as differentially expressed for an absolute log2 fold change (‘beta’ coefficient) ≥ 1 and a q-value ≤ 0.01.

#### Alternative splicing Events

For detecting alternative splicing events, the SUPPA2 algorithm (version 2.3) was used (Trincado et al., 2018). Kallisto transcript quantification (transcripts per million) values for all samples and GTF file of the assembled transcriptome were used as input. Subsequent Python scripts were executed according to the provided guidelines.

#### Differential transcript usage

Differential transcript usage (DTU) was analyzed by using Kallisto transcripts quantifications (estimated counts rounded to integer) as counting bins for fitting transcript-level negative binomial models in DEXSeq (Anders et al., 2012;Soneson et al., 2016). These models were fitted for pairwise comparisons of interest and transcripts were required to have at least 10 counts for three out of six samples. Hence, instead of differential exon usage, DEXSeq will test significant DTU in *sme1-2*. DTU was assessed for 14,278 multi-transcript genes encoding 52,618 transcripts. Subsequently, stageR was used for DTU analysis as described (Van den Berge et al., 2017). In the first stage, *i*.*e*. screening stage, transcript *P*-values were aggregated to the gene level and a 5% overall FDR level was used as threshold. For the significant genes, transcripts were confirmed in the second stage.

### Gene set enrichment analysis

Differential gene or protein lists were used for gene set overrepresentation analysis using the online webtool DAVID (Huang da et al., 2009). Gene sets included gene ontology (GO) biological process, KEGG pathways and SMART protein domains.

## SUPPLEMENTAL INFORMATION

The following materials are available in the online version of this article.

### Supplemental Materials and methods

**Supplemental Figure 1.** Nonsense mutation *40*.*2* in *SME1* (AT2G18740).

**Supplemental Figure 2.** Plant growth under chemical and genetic perturbations of the spliceosome.

**Supplemental Figure 3.** Growth-related abiotic stress assays on *SME2* knockdown mutant plants

**Supplemental Figure 4.** Normalized log2 peptide intensity plots (∼protein intensity) for anthocyanin biosynthesis proteins (**A**) TRANSPARENT TETA5 (TT5, AT3G55120) and (**B**) UDP-GLUCOSE: ANTHOCYANIDIN 3-O-GLUCOSYLTRANSFERASE (UF3GT, AT5G54060), as obtained by MSqRob (Goeminne et al., 2016)/

**Supplemental Figure 5.** Nonsense-mediated decay (NMD) hallmarks in Araport11 and Scallop-derived transcripts.

**Supplemental Figure 6.** Snurportin-1 (SPN1) AT4G24880 (**A**) and DHX35 AT4G18465 (**B**) alignments with orthologs in human, mouse, Drosophila, Physcomitrium (moss), and wheat.

**Supplemental Figure 7.** AT5G13310 and AT5G13970 as putative TSSC4 ortologs.

**Supplemental Figure 8.** Rosette area of wild-type (black) and mutant (orange) *spn1, icln-2, dhx35, tssc4a, and tssc4b* seedlings (n = 6, mean ± 95% confidence interval) grown on horizontal plates with control media or media supplemented with 25 nM methyl viologen (MV), 25 mM mannitol, or 50 mM NaCl.

**Supplemental Figure 9.** Quantitative real-time PCR showing the relative expression for U snRNAs in wild-type plants (grey) and mutants (orange) for putative pre-mRNA splicing proteins SPN1 (AT4G24880, mutant SALK_119615.9).

**Supplemental Figure 10.** Genome read coverage maps for AS genes in wild-type (black) and *sme1-2* (orange) for (A) *ACONITASE1* (*ACO1*, AT4G35830), (B) *COPPER/ZINC SUPEROXIDE DISMUTASE1* (*CSD1*, AT1G08830) and (C) *CIRCADIAN CLOCK ASSOCIATED1* (*CCA1*, AT2G46830).

**Supplemental Table 1.** Differential mRNA-seq analysis by the kallisto-sleuth pipeline (see Methods).

**Supplemental Table 2.** Differential protein abundance by MaxQuant-MSqRob processing (see Methods).

**Supplemental Table 3.** Gene set enrichments for differentially expressed genes and proteins in *sme1-2*.

**Supplemental Table 4.** SUPPA2 (Trincado et al., 2018) results for AS events in *sme1-2* versus wild-type.

**Supplemental Table 5.** Differential transcript usage (DTU) analysis of *sme1-2* versus wild-type.

**Supplemental Table 6.** Gene set enrichments for the 425 AS genes identified by both the SUPPA2 and DTU analysis.

**Supplemental Table 7.** Identified protein interactors of SME1. Interactors are classified according to (Koncz et al., 2012). In addition, GEMIN2 (Schlaen et al., 2015) and ICLN (Huang et al., 2016) act as spliceosome assembly proteins in Arabidopsis.

**Supplemental Table 8.** Primers used for genotyping.

**Supplemental Table 9.** Primers used for qPCR. UBC10 and PP2AA3 were reference genes.

**Supplemental Table 10.** Aligned reads per sample. Statistics were extracted from the Kallisto mapping logs.

## Acknowledgements

We are grateful to Veronique Storme for statistical analysis, Koen Van den Berge for helping with the AS analysis, and Martine De Cock for help in preparing the manuscript.

## Funding

This work was supported by the Research Foundation-Flanders–Fonds de la Recherche Scientifique (FWO-FNRS The Excellence of Science [EOS] Research project no. 30829584 and FWO project no. G055416N to F.V.B.), the Agency for Innovation by Science and Technology (Industrial R&D project no. 100555 and Phoenix project 070347), Ghent University (‘Bijzonder Onderzoeks Fonds’ project no. 01J11311). P.W. is the recipient of a Junior Postdoctoral fellowship (project no. 12T1722N) of the Research Foundation-Flanders for and V.V.R. was indebted to the Agency for Innovation by Science and Technology for a predoctoral fellowship (project no. 141029).

## Conflict of interest

The authors declare that they have no conflict of interest.

